# Google Trends and Online Q&A site Reveal Surging Demand for Endemic Pet Reptiles in Japan

**DOI:** 10.1101/2024.04.28.591490

**Authors:** Richard Digirolamo

## Abstract

The global trade in pet reptiles is substantial, with Japan being a major importer. Understanding pet reptile preferences is crucial for conservation, as overexploitation of wild populations can occur. This study aimed to determine Japan’s most popular pet reptiles, including traded and endemic species, using the largest question-answering website in Japan, Yahoo! Chiebukuro and Google Trends as data sources. We analysed data from 2004 to 2023, focusing on recent periods (2020-2023) for a comprehensive assessment of public interest in 20 different pet reptile species. Data revealed a significant overlap between the two tools in identifying popular species. Leopard geckos were consistently the most popular, aligning with trade data. Surprisingly, the endemic Japanese grass lizard surpassed the popularity of traditionally favoured species like central bearded dragons. A concerning preference for invasive turtles (Chinese pond turtles, red-eared sliders) was also noted. These findings highlight the popularity of both traded and readily available endemic species within Japan’s pet market. This emphasises the need to monitor domestic trends alongside the traditional trade focus to ensure the sustainability of wild reptile populations. Yahoo! Chiebukuro and Google Trends offer valuable tools for such monitoring. Google Trends’ regional data pinpointed Okinawa prefecture’s high interest in leopard geckos, raising concerns about potential invasiveness. This study demonstrates the importance of a comprehensive approach to understanding pet reptile popularity. The combined use of these tools and the adaptable methodology provide a template for monitoring trends in Japan and other countries, contributing to proactive conservation efforts within the global pet trade.

## Introduction

The global trade in non-avian reptiles (hereafter reptiles) as exotic pets represents a significant market, with imports exceeding $61 million in 2022. Japan is a major driver of this demand, having 20% of the global reptile imports share in 2022 (Trendeconomy, 2023). In Japan, reptiles are consistently the most imported animal group, including illegal imports (Kitade & Naruse, 2020). The high demand for pet reptiles, fueled in part by their attractiveness and perceived novelty, can lead to the overexploitation of wild populations, particularly for rare and sought-after species (Marshall et al., 2020; Courchamp et al., 2006; Wakao et al., 2018; Kitade & Naruse, 2020).

While the global reptile trade involves thousands of species, commonly available pet reptiles tend to be widespread, captive-bred, and/or domestically obtained. Due to their generally non-threatened status, these species are often unregulated, leading to a vast and poorly monitored segment of the pet trade (Valdez, 2021). This may be particularly true in Japan, where endemic species such as the Japanese pond turtle (*Mauremys japonica*), Japanese skink (*Plestiodon japonicus*), Japanese grass lizard (*Takydromus tachydromoides*), Schlegel’s Japanese gecko (*Gekko japonicus*), and Japanese rat snake (*Elaphe climacophora*) are taken from the wild as pets. Anecdotal evidence suggests their suitability as pets is promoted online, along with methods for their capture. The same is mentioned in local literature (Kawazoe, 2020; Takenaka, 2023; Nieve & Wildsky, 2023; Tsuruno, 2003; REP FAN Vol.20, 2023). Real-time monitoring of these species’ popularity as pets is crucial for conservation efforts, even if they are not currently endangered.

Google Trends, a tool analysing top Google Search queries, offers a potential method for real-time monitoring of pet reptile preferences. It provides access to a largely unfiltered sample of search requests, offering insights into public interest across various regions and languages (FAQ About Google Trends Data - Trends Help, n.d.; Rogers, 2016). Google Trends has been applied in diverse fields, including medicine (Pelat et al., 2009; Bragazzi & Mahroum, 2019; Lippi et al., 2022; Díaz et al., 2023), economics (Choi & Varian, 2012; Bantis et al., 2023), and conservation biology (Zieger & Springer 2020, 2021). Its recent use in assessing public interest in vertebrate species (Holmes et al., 2022) and traded pet reptiles globally (Valdez, 2021) demonstrates its value in understanding wildlife-related trends.

Valdez (2021) highlighted the potential of Google Trends in identifying popular pet reptiles while demonstrating discrepancies with traditional survey methods. Surveys are often limited by sample size and potential for bias (Butler, 2013), whereas Google Trends may provide a more robust assessment of public interest (Vosen & Schmidt, 2011). However, despite Japan’s substantial reptile imports, Valdez (2021) found no data for Japan in its current popularity search results using Google Trends. The study suggests that this is likely due to insufficient search volumes. This is unexpected given Japan’s large pet reptile market (Trendeconomy, 2023; Wakao et al., 2018) and >80% of the population’s internet usage since 2013, and Google is the most used search engine with its share of >70% (Ministry of Internal Affairs and Communications, 2021, 2022). This discrepancy may stem from Google Trends limitations in search terminology, as Google Trends can be sensitive to non-English language (Mavragani et al., 2018).

Yahoo! Chiebukuro, Japan’s largest question-answering website (National Institute of Informatics, 2024), offers another potential measure of pet reptile popularity. Users post questions on various topics and manually categorise them for ease of browsing. Theoretically, higher interest in a topic should correlate with a more significant number of associated questions. This has been demonstrated in studies on drug abuse trends (Kariya et al., 2023).

This study aims to determine Japan’s most popular pet reptiles, including traded and endemic species, using Yahoo! Chiebukuro and Google Trends (with refined terminology) as data sources. Data from 2004-2023 will be analysed, focusing on recent trends (2020-2023) and comparing the two data sources.

## Methods

### Listing of Popular Pet Reptiles

Twenty popular pet reptile species were selected based on their prevalence in the global pet trade (Valdez, 2021), representation in Japanese pet-focused publications and websites, and the authors’ expertise. To align with Japanese regulations, we excluded Boa constrictor (*Boa constrictor*) and Reticulated python (*Malayopython reticulatus*). While popular globally (Valdez, 2021), these species are prohibited as pets in Japan (Ministry of the Environment, 2019). The final list is provided in Table 1.

**TABLE 1.**
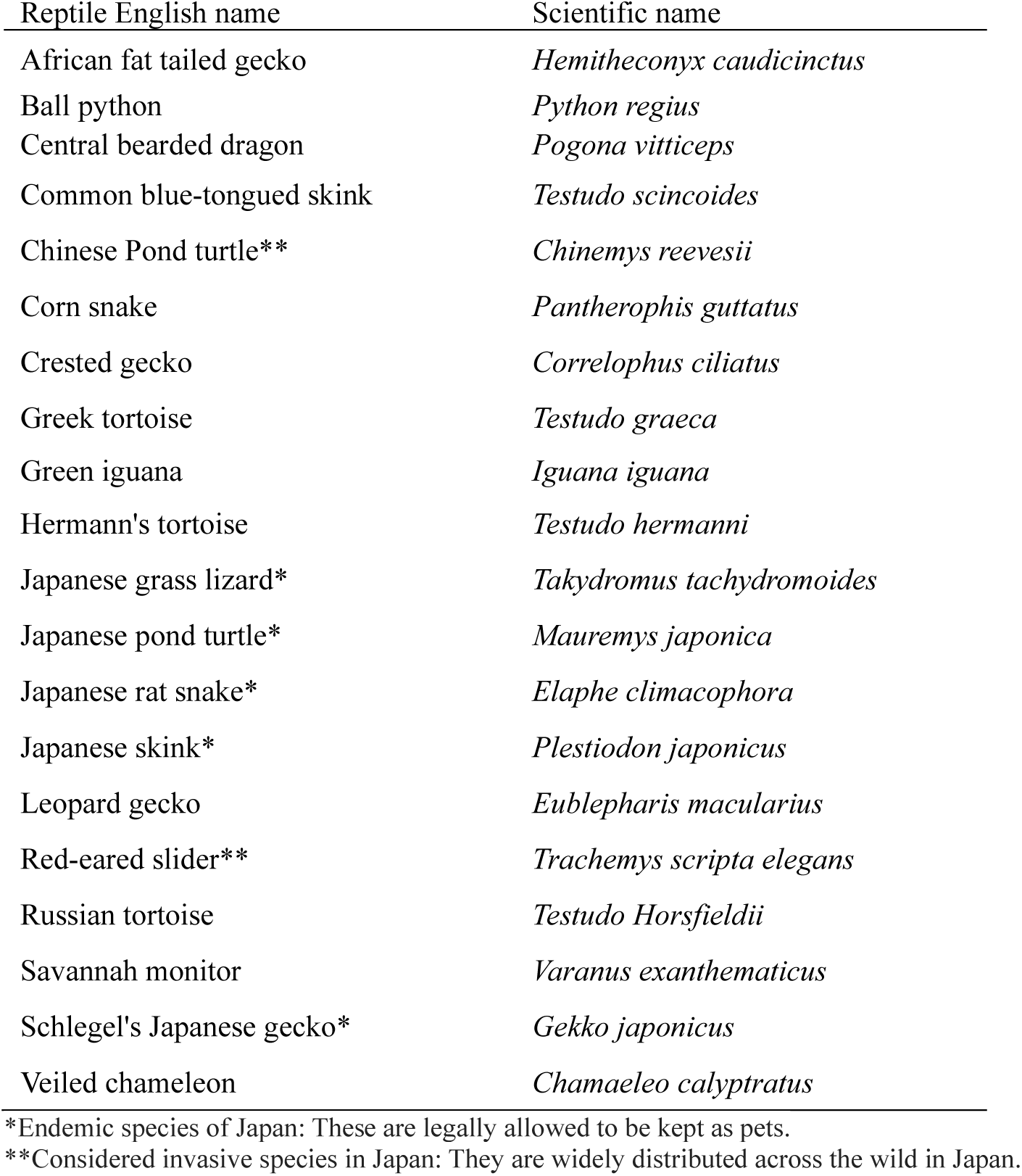
List of the 20 selected reptile species (alphabetical order).

### Yahoo! Chiebukuro Data

Yahoo! Chiebukuro (https://chiebukuro.yahoo.co.jp/), Japan’s largest question-and-answer platform, was used as a data source. Each species from Table 1 was searched using its Japanese name(s) within the “Pet” category. Search settings were iterated to cover yearly periods from 2004 to 2023. The total number of questions per year was recorded and divided into three periods: 1) Grand Total (2004-2023): Overall popularity across the entire period. 2) Past (2004-2019): Pre-COVID-19 pandemic trends, excluding the COVID-19 pandemic pet ownership surge (Rakuten Insight, Inc. (2021). 3) Recent (2020-2023): Recent popularity, potentially influenced by the pandemic. To ensure comprehensive data capture, searches included common Japanese aliases for each species (see Supplementary Material Table 1). Data collection occurred in March 2024.

### Google Trends Data

Google Trends (https://trends.google.com/trends/), a freely available tool that analyses search query patterns within Google Search, was used to assess online search popularity. Due to Google Trends’ five-item comparison limit, reptile search trends were analysed in groups. The most popular reptile was established as a benchmark and included in each group alongside four other reptiles, allowing for a comparative analysis of relative temporal trends. Since Google Trends provides relative search frequencies, results are sensitive to the benchmark term (Nghiem et al., 2016). A yearly moving average was applied to mitigate noise and emphasise long-term trends. This approach also reduced the impact of seasonal fluctuations, accounting for higher summer search volumes for reptiles (Valdez, 2021). Each species from Table 1 was typed in Japanese and searched with its auto-translated English name, which appears in the search field, with the following parameters: a) Period: Yearly, 2004-2023 b) Region: Japan c) Search Topic: “Reptiles” or “Snake” (for snake species) d) Category: “Pets”. Yearly average search volume indexes were split into “Total Average (2004-2023)”, “Past Average (2004-2019)”, and “Recent Average (2020-2023)”, aligning with the Yahoo! Chiebukuro analysis. Standard deviations were calculated. Data collection occurred in March 2024. Specific search terms are detailed in Supplementary Material Table 2.

## Results

### Yahoo! Chiebukuro

Fig 1a shows the top five reptile species’ highest number of questions generated in Yahoo! Chiebukuro. Analysis of Yahoo! Chiebukuro data revealed the highest overall question volume for leopard geckos. Focusing on the recent period (2020-2023), leopard geckos remained dominant (11,449 questions), followed by Japanese grass lizards (2760 questions), central bearded dragons (2251 questions), Chinese pond turtles (2184 questions), and red-eared sliders (1847 questions).

**FIG. 1.**
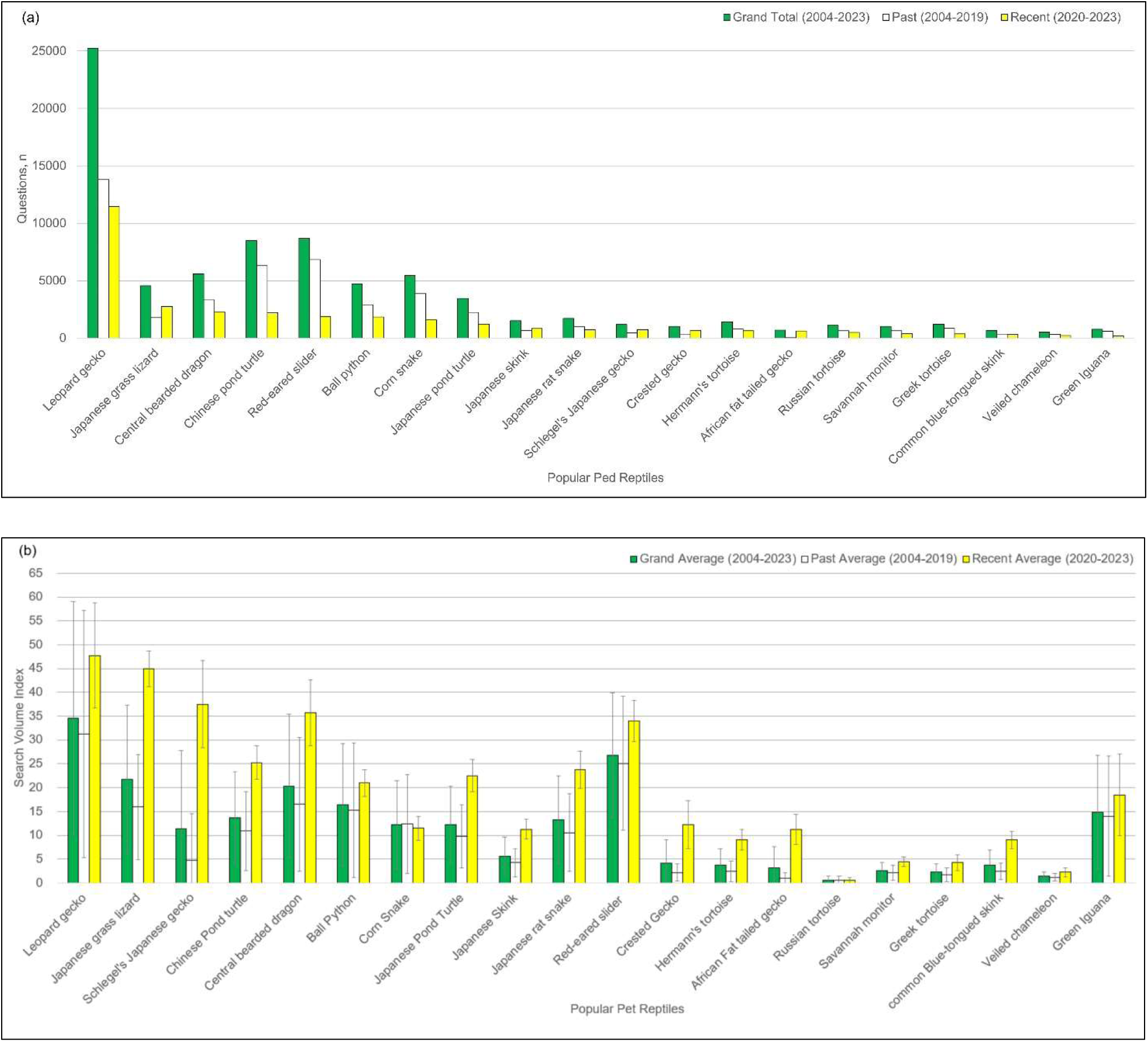
(a) The total number of questions raised in Yahoo! Chiebukuro per reptile species is divided into periods: past (2004-2019), recent (2020-2023), and their total. (b) The average search volume index per reptile species on Google Trends is divided into periods in the same way as (a).

### Google Trends

Fig 1b shows the top five reptile species’ highest average search volume index (SVI) on Google Trends. Focusing on the recent period (2020-2023), leopard geckos exhibited the highest score (SVI 48, SD=11.00). They were closely followed by Japanese grass lizards (SVI 45, SD=3.74), Schlegel’s Japanese geckos (SVI 38, SD=9.11), central bearded dragons (SVI 36, SD=6.95), and red-eared sliders (SVI 34, SD=4.32).

### Trend Analysis (2004-2023)

Fig 2a shows trends of the number of questions from 2004-2023 for the top five reptile species in Yahoo! Chiebukuro. Leopard geckos consistently ranked first since 2012. Interest in red-eared sliders and Chinese pond turtles declined after 2011 and 2015, respectively. Central bearded dragons rose in popularity and were the second-highest species in 2017 but trended closely with Chinese pond turtle since then. Japanese grass lizards saw a dramatic three-times surge in 2021 compared to 2019 and continued its surge as of 2023.

**FIG. 2.**
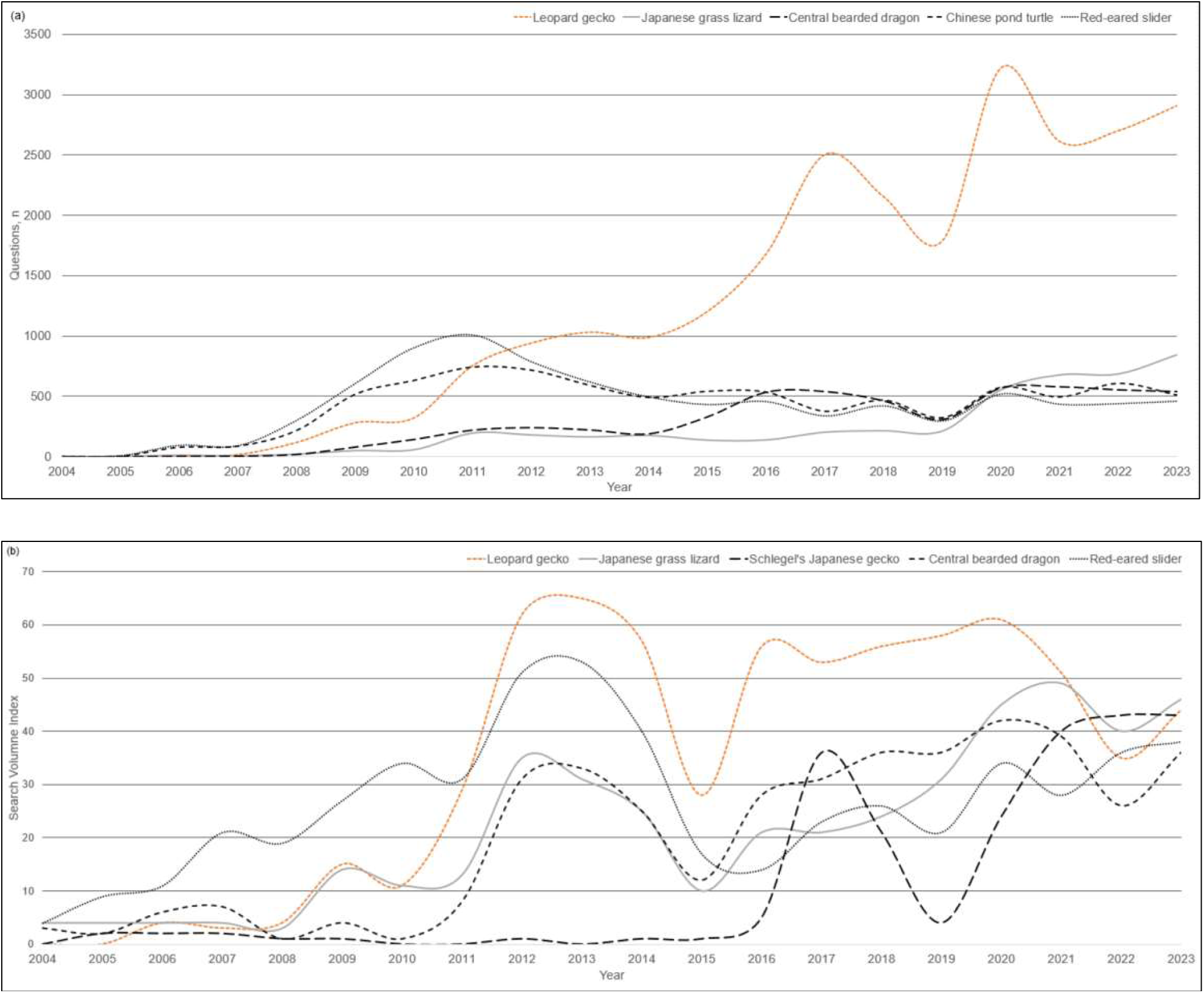
(a) Yahoo! Chiebukuro trends of number of questions raised from 2004-2023 for the top five species. (b) SVI on Google Trends from 2004-2023 for the top five species. The number represents the yearly average.

Google Trends (Fig. 2b) indicated a 2012-2013 popularity spike for most top-ranked species, followed by fluctuations. Schlegel’s Japanese geckos exhibited a distinct trend, peaking in 2017, followed by a sharp drop until 2019, then another rise until 2022, and a stay level in 2023.

### Subregional Analysis

Table 2 presents Google Trends data on interests by subregions (top three prefectures) for the five most popular species from Google Trends. Notable findings are that Okinawa prefecture, Japan’s most southern sub-tropical island region, had the highest search interests for central bearded dragons and third highest with margins of one point difference from first and second prefecture for leopard geckos.

**TABLE 2.** Regional interest in the top five reptile species as measured by Google Trends Search Volume Index (SVI).

| Species | 1st Prefecture (SVI) | 2nd Prefecture (SVI) | 3rd Prefecture (SVI) |
| --- | --- | --- | --- |
| Leopard gecko | Wakayama (100) | Gifu (100) | Okinawa (99) |
| Japanese grass lizard | Miyagi (100) | Iwate (84) | Akita (75) |
| Schlegel's Japanese gecko | Kagoshima (100) | Shizuoka (90) | Miyazaki (87) |
| Central bearded dragon | Okinawa (100) | Kochi (79) | Yamanashi (77) |
| Red-eared slider | Okayama (100) | Tokushima (91) | Wakayama (80) |

### Overall Popularity Ranking

Table 3 compares the popularity of the reptile species based on results from Yahoo! Chiebukuro and Google Trends searches. The data indicates that both platforms show the leopard gecko as the most popular reptile of those analysed. The Japanese grass lizard, central bearded dragon, and red-eared slider are the top five most popular species across both platforms. Of the top ten ranked species, Chinese pond turtle, ball python, corn snake, and Japanese skink see higher interest in Yahoo! Chiebukuro than their Google Trends ranking. Conversely, Schlegel’s Japanese gecko, Japanese rat snake and green iguana rank significantly higher on Google Trends than its Yahoo! Chiebukuro position.

**TABLE 3.** Popularity of reptile species on Yahoo! Chiebukuro (Yahoo) and Google Trends (Google). Yahoo represents the total number of questions generated per species from 2020-2023, while Google represents the average search volume index (SVI) for the same period.

| Species | Rank<br>(Yahoo / Google) | Yahoo<br>Question, n (2020-2023) | Google<br>SVI (2020-2023) |
| --- | --- | --- | --- |
| Leopard gecko | 1/1 | 11449 | 48 |
| Japanese grass lizard | 2/2 | 2760 | 45 |
| Central bearded dragon | 3/4 | 2251 | 36 |
| Chinese pond turtle | 4/6 | 2184 | 25 |
| Red-eared slider | 5/5 | 1847 | 34 |
| Ball python | 6/9 | 1820 | 21 |
| Corn snake | 7/11 | 1578 | 12 |
| Japanese pond turtle | 8/8 | 1218 | 23 |
| Japanese skink | 9/13 | 820 | 11 |
| Japanese rat snake | 10/7 | 716 | 24 |
| Schlegel's Japanese gecko | 11/3 | 712 | 38 |
| Crested gecko | 12/11 | 668 | 12 |
| Hermann's tortoise | 13/15 | 635 | 9 |
| African fat-tailed gecko | 14/13 | 616 | 11 |
| Russian tortoise | 15/20 | 482 | 1 |
| Savannah monitor | 16/17 | 375 | 5 |
| Greek tortoise | 17/18 | 352 | 4 |
| Common blue-tongued skink | 18/15 | 317 | 9 |
| Veiled chameleon | 19/19 | 203 | 2 |
| Green iguana | 20/10 | 179 | 19 |

## Discussion

### Consistency of Findings & Implications for Reptile Popularity Tracking

This study demonstrated the first quantitative assessment of pet reptile popularity in Japan, employing Yahoo! Chiebukuro and Google Trends. Yahoo! Chiebukuro and Google Trends significantly overlapped in identifying popular pet reptiles in Japan. Eight species appeared within the top ten rankings of both sources. This consistency suggests that both tools hold the potential for real-time monitoring of pet reptile preferences, offering potential advantages over traditional survey methods, which can be limited by sample size, bias, and time consumption.

The prominence of five species found in the wild (Japanese grass lizard, Chinese pond turtle, red-eared slider, Japanese pond turtle, and Japanese rat snake) in the top ten rankings of both sources highlights the substantial demand within the Japanese reptile pet market. Three of these are endemic species (Japanese grass lizard, Japanese pond turtle and Japanese rat snake) are of conservation concern, with the Japanese grass lizard listed as “Vulnerable” and “Near Threatened” in two prefectures, Japanese pond turtle listed as “Endangered” and “Near Threatened” in 29 prefectures, and Japanese rate snake listed as “Near Threatened” in four prefectures (Search System of Japanese Red Data, 2020). Japanese pond turtles, in particular, are showing significant endangerment in many prefectures, and pet dealers’ overexploitation is considered one of the reasons (Yasukawa et al., 2008). This raises questions about the sustainability of continuously harvesting wild populations for the pet and the potential long-term impacts on these species’ viability.

### Leopard Gecko: The Most Popular Pet Reptile

Leopard geckos were unequivocally the most popular pet reptile in Japan. This aligns with trade data (Wakao et al., 2018). Several factors likely contribute to their dominance, including a) manageability: Small size, nocturnal habits, and relatively low space requirements b) ease of care: Leopard geckos are considered hardy with straightforward husbandry needs and c) affordability: They are readily available all year round through captive breeding, keeping prices relatively low for common morphs (Terao, 2021; Nakagawa, 2021).

While Yahoo! Chiebukuro data shows a sustained and overwhelming interest in leopard geckos, Google Trends suggests a possible decrease in their relative popularity compared to other species. This discrepancy highlights how Google Trends scores are calculated based on relative search volumes. A decline in their score does not necessarily indicate decreased interest but may reflect the rising popularity of other species.

### Invasive Turtles: A Concerning Preference

The popularity of invasive turtle species, particularly the Chinese pond turtle and red-eared slider, is a cause for concern. Despite the Japanese pond turtle’s historical prominence in the pet trade (Wakao et al., 2018), recent wild population surveys indicate that invasive turtles significantly outnumber native species (The Nature Conservation Society of Japan, 2023; Yabe, 2014). Several factors may explain this shift including a) accessibility: wild-caught invasive turtles may be readily obtained, unlike rare Japanese pond turtles b) cost: Chinese pond turtles are generally less expensive than their native counterparts in trade c) legal status: while red-eared sliders are now banned from trade in Japan, existing pets can still be kept, leading to a potential surplus available for free adoption (*Red-eared Slider | Countermeasures Against Invasive Species in Japan | Invasive Species Law*, 2023). This preference for invasive turtles poses significant ecological risks. Both species can outcompete native turtles for resources, and due to their longevity, irresponsible pet owners may unlawfully release them into the wild. Both species originated from irresponsible pet release and continuation of it at present, leading to the current population dominance (The Nature Conservation Society of Japan, 2023; Yabe, 2014, *Red-eared Slider | Countermeasures Against Invasive Species in Japan | Invasive Species Law*, 2023). The continued demand for these species as pets could exacerbate existing invasion problems and threaten native turtle populations.

### Popularity of Endemic Species: Drivers and Implications

The unexpected popularity of Japanese grass lizards and Japanese rat snakes, both rarely commercially traded, raises intriguing questions. Their appeal likely stems from a) ease of acquisition: As wild-caught species, they are obtainable without cost. b) small size (Japanese grass lizard): Japanese grass lizard is small (adult snout-vent length 5-6 cm) (Takenaka, 1981, 2023), which makes them suitable for smaller living spaces and less demanding in terms of enclosure setup. c) cultural significance (Japanese rat snake): Historically considered symbols of good fortune (Digital Museum of Hiroshima University, 2024), rat snakes may benefit from positive cultural associations.

The popularity of these endemic species underscores the need for careful monitoring of wild populations. While currently not considered endangered (Kidera & Ota 2017a, 2017c), unregulated harvesting for pets could potentially lead to local population declines, particularly as both species have existing regional conservation concerns.

### Shifting Preferences: Smaller and Local

While still popular, central bearded dragons have shown signs of levelling off since 2017-2018, overtaken by smaller, endemic reptiles. This suggests a potential preference shift towards even more manageable pets, particularly those readily sourced locally.

### Conservation Value of Google Trends’ Regional Data

Google Trends’ ability to pinpoint regional interests is a valuable tool for conservation. The identified high interest in leopard geckos within Okinawa prefecture raises specific concerns. Okinawa’s subtropical climate resembles some of the leopard gecko’s native habitats (Khan, 2009, *Climate of Okinawa Prefecture*. (n.d.)), increasing the risk of establishment if pets are irresponsibly released. This could have devastating consequences for smaller endemic ground-dwelling geckos, the *Goniurosaurus kuroiwae*, which includes several sub-species and are rated between endangered and vulnerable (Kidera & Ota, 2017b; Toda, 2024). Targeted monitoring, educational campaigns, and stricter regulations on leopard gecko ownership in Okinawa may be necessary to mitigate this risk. Central bearded dragons, on the other hand, also showed high interest in Okinawa prefecture, the sub-tropical climate with more than twice as much precipitation as its arid/semi-arid natural habitat in Australia (*Climate of Okinawa Prefecture*. (n.d.). Australian Government Department of Climate Change, Energy, the Environment and Water, 2021) is unlikely to meet their desired environment to thrive. However, captive-bred reptiles locally, for several generations, may be able to adapt to the local climate; thus, the potential for invasiveness should be considered, and monitoring should still be conducted.

### Limitations

Yahoo! Chiebukuro and Google Trends proved valuable for assessing pet reptile popularity, demonstrating consistency with additional insights. Both have limitations:

a. Exclusion of non-users: Results may not reflect populations who do not use these tools.
b. Search term sensitivity: Accurate results depend on using all common names and aliases for Yahoo! Chiebukuro. If any questions are posted using uncommon names or abbreviations, they will not be picked up in the search results. Google Trends’ auto-translation to a single name available to choose under a reptile topic can introduce bias for species with several common names. For example, the Chinese pond turtle is also known as Reeve’s Pond turtle. When the common turtle’s name was entered in the search term in Japanese, it automatically displayed “Chinese pond turtle” under the topic “Reptile”. This assumes Google collects all Japanese searches for the turtle under the Chinese pond turtle. A search using the English name “Reeve’s Pond turtle” by English language entry generated fewer values than “Chinese pond turtle” (data not shown). This showed that it is important to type the name of a local language species into the search term and select the correct auto-translated name within Google Trends.
c. Categorisation errors: Yahoo! Chiebukuro relies on user-selected categories, poses unlikely but possible mis-selection, e.g., instead of asking under the pet category, they may select the category Biology/Animals. For Google Trends the exact mechanism behind Google Trends’ topic categorisation is unknown. Google Trends’ lack of transparency regarding its categorisation mechanism may pose challenges for accurately interpreting pet reptile popularity data. A search that includes both a species name and pet care terms (e.g., “how to care for leopard gecko”) might be correctly categorised by Google. However, a simple species search like “leopard gecko,” followed by accessing a website containing pet and scientific information, creates ambiguity. Google may struggle to determine the user’s intent (pet-related vs. general scientific curiosity).

These limitations could explain discrepancies between Yahoo! Chiebukuro and Google Trends results for species like the green iguana and Schlegel’s Japanese gecko. Due to the species’ specialised needs, Green iguana owners may use more targeted searches from Google than Yahoo! Chiebukuro. In contrast, the house-dwelling Schlegel’s Japanese geckos, commonly seen in Japan (Wada, 2003), might generate more general scientific inquiries alongside pet-related searches. For example, the highly accessed website Wikipedia containing mixed content (pet care and biology) (Schlegel’s Japanese Gecko, 2024) can complicate the categorisation issue. Without understanding Google Trends’ categorisation process, conclusions about pet popularity must be drawn cautiously, particularly for abundant species such as Schlegel’s Japanese gecko.

## Conclusions

This study provides the first quantitative assessment of pet reptile popularity in Japan using Yahoo! Chiebukuro and Google Trends. A key finding was the significant popularity of endemic, non-traded species alongside traditional pet reptiles. This highlights the importance of monitoring trade-driven and domestic pet preference trends for effective conservation efforts. The surprising popularity of Japanese grass lizards, surpassing established favourites like central bearded dragons and ball pythons, underscores the dynamic nature of the pet market. Leopard geckos’ dominance as the most popular pet reptile aligns with existing trade data.

Both research tools demonstrated strengths and limitations, suggesting their combined use is the most effective. Google Trends’ ability to pinpoint regional interests, such as the high interest in leopard geckos within Okinawa prefecture, is invaluable for targeted monitoring of potential invasive species threats. However, its reliance on relative search volumes can sometimes obscure absolute popularity trends, as seen with leopard geckos compared to Japanese grass lizards. Yahoo! Chiebukuro provided crucial context by revealing absolute question counts within the pet category. While Google Trends’ translated terms accuracy in aggregating aliases and categorisation remain potential concerns, Yahoo! Chiebukuro offers a more controlled data source in this regard.

Smaller reptiles’ increasing popularity as traded and endemic pets emphasises the need for comprehensive monitoring. Understanding these trends is crucial to predicting future pet trade demands and proactively addressing conservation concerns. Japan’s history of overexploitation leading to endangerment (e.g., *Goniurosaurus kuroiwae*) (Toda, 2024) underscores this urgency. Yahoo! Chiebukuro and Google Trends, used in conjunction, provide a valuable toolkit for tracking pet reptile popularity and potentially for other animal groups. The methodology developed in this study offers a template adaptable to other countries and languages, facilitating broader monitoring of endemic species’ popularity within the pet trade.

## Acknowledgements

This research received no specific grant from any funding agency or commercial or not-for-profit sectors.

## Ethical standards

None.

## Data availability

Yahoo! Chiebukuro and Google Trends data are available from respective websites as described in the methods section.

