## Supplemental Material for "Google Trends and Online Q&A site Reveal Surging Demand for Endemic Pet Reptiles in Japan"

RICHARD DIGIROLAMO

SUPPLEMENTARY TABLE 1 List of 20 Reptile Species with Common Names and Japanese Terms. This table provides a comprehensive list of 20 reptile species commonly kept as pets, along with their English common names, scientific names, and corresponding Japanese terms used for searching on Yahoo! Chiebukuro. Additionally, alternative search terms for each species are included in the table third column to ensure comprehensive search results.

| Reptile English name (Other common names) | Scientific name | Japanese Name(s) searched in Yahoo! Chiebukuro |
|---|---|---|
| African fat tailed gecko | *Hemitheconyx caudicinctus* | ニシアフリカトカゲモドキ |
| Ball python | *Python regius* | ボールパイソン<br>ボールニシキヘビ |
| Central bearded dragon | *Pogona vitticeps* | フトアゴヒゲトカゲ |
| Common blue-tongued skink | *Testudo scincoides* | アオジタトカゲ |
| Chinese Pond turtle (Reeve's pond turtle) | *Chinemys reevesii* | クサガメ<br>ゼニガメ |
| Corn snake | *Pantherophis guttatus* | コーンスネーク |
| Crested gecko | *Correlophus ciliatus* | オウカンミカドヤモリ<br>クレステッドゲッコー |
| Greek tortoise | *Testudo graeca* | ギリシャリクガメ |
| Green iguana | *Iguana iguana* | グリーンイグアナ |
| Hermann's tortoise | *Testudo hermanni* | ヘルマンリクガメ |
| Japanese grass lizard | *Takydromus tachydromoides* | ニホンカナヘビ<br>カナヘビ |
| Japanese pond turtle | *Mauremys japonica* | ニホンイシガメ |
| Japanese rat snake | *Elaphe climacophora* | アオダイショウ |
| Japanese skink (Japanese five-lined skink) | *Plestiodon japonicus* | ニホントカゲ |
| Leopard gecko | *Eublepharis macularius* | ヒョウモントカゲモドキ<br>レオパ |
| Red-eared slider | *Trachemys scripta elegans* | ミシシッピーアカミミガメ<br>ミドリガメ |
| Russian tortoise | *Testudo Horsfieldii* | ロシアリクガメ<br>ヨツユビリクガメ |
| Savannah monitor | *Varanus exanthematicus* | サバンナモニター |
| Schlegel's Japanese gecko (Japanese gecko) | *Gekko japonicus* | ニホンヤモリ |
| Veiled chameleon | *Chamaeleo calyptratus* | エボシカメレオン |

SUPPLEMENTARY TABLE 2 This table provides a comprehensive list of 20 reptile species commonly kept as pets, along with their scientific names, English common names, corresponding Japanese names used for searching on Google Trends, and the auto-translated names generated by Google Trends. The auto-translated names are categorized into either "Reptile" or "Snake" topics.

| Reptile English name (Other common names) | Scientific name | Japanese names entered in Google Trends | Google Trends auto translated names |
|---|---|---|---|
| African fat-tailed gecko | *Hemitheconyx caudicinctus* | ニシアフリカトカゲモドキ | African fat-tailed gecko |
| Ball python | *Python regius* | ボールパイソン | Ball python |
| Central bearded dragon | *Pogona vitticeps* | フトアゴヒゲトカゲ | Central bearded dragon |
| Common blue-tongued skink | *Testudo scincoides* | アオジタトカゲ | Common blue-tongued skink |
| Chinese Pond turtle (Reeve's pond turtle) | *Chinemys reevesii* | クサガメ | Chinese pond turtle |
| Corn snake | *Pantherophis guttatus* | コーンスネーク | Corn snake |
| Crested gecko | *Correlophus ciliatus* | オウカンミカドヤモリ | Crested gecko |
| Greek tortoise | *Testudo graeca* | ギリシャリクガメ | Greek tortoise |
| Green iguana | *Iguana iguana* | グリーンイグアナ | Green iguana |
| Hermann's tortoise | *Testudo hermanni* | ヘルマンリクガメ | Hermann's tortoise |
| Japanese grass lizard | *Takydromus tachydromoides* | ニホンカナヘビ | Takydromus tachydromoides |
| Japanese pond turtle | *Mauremys japonica* | ニホンイシガメ | Japanese pond turtle |
| Japanese rat snake | *Elaphe climacophora* | アオダイショウ | Japanese rat snake |
| Japanese skink (Japanese five-lined skink) | *Plestiodon japonicus* | ニホントカゲ | Japanese skink |
| Leopard gecko | *Eublepharis macularius* | ヒョウモントカゲモドキ | Leopard gecko |
| Red-eared slider | *Trachemys scripta elegans* | ミドリガメ | Pond slider |
| Russian tortoise | *Testudo Horsfieldii* | ヨツユビリクガメ | Russian tortoise |
| Savannah monitor | *Varanus exanthematicus* | サバンナモニター | Savannah monitor |
| Schlegel's Japanese gecko (Japanese gecko) | *Gekko japonicus* | ニホンヤモリ | Schlegel's Japanese gecko |
| Veiled chameleon | *Chamaeleo calyptratus* | エボシカメレオン | Veiled chameleon |
